# Cingulate and striatal hubs are linked to early skill learning

**DOI:** 10.1101/2024.11.20.624544

**Authors:** Hisato Sugata, Fumiaki Iwane, William Hayward, Valentina Azzollini, Debadatta Dash, Roberto F Salamanca-Giron, Marlene Bönstrup, Ethan R Buch, Leonardo G Cohen

## Abstract

Early skill learning develops in the context of activity changes in distributed cortico-subcortical regions. Here, we investigated network hubs—centers of information integration and transmission—within the brain network supporting early skill learning. We recorded magnetoencephalographic (MEG) brain activity in healthy human subjects who learned a moderately difficult sequence skill with their non-dominant left hand. We then computed network hub strength by summing top 10% functional connectivity over 86 parcellated brain regions (AAL3 atlas) and five brain oscillatory frequency bands (alpha, low-, high-beta, low- and high-gamma). Virtually all skill gains developed during rest intervals of early learning (micro-offline gains). MEG hub strength in the alpha band (8-13Hz) in bilateral anterior cingulate (ACC) and caudate and in the low-beta band (13-16Hz) in bilateral caudate and right putamen correlated with micro-offline gains. These regions linked strongly with the hippocampus, parahippocampal cortex, and lingual and fusiform gyri. Thus, alpha and low-beta brain oscillatory activity in cingulate and striatal regions appear to contribute as hubs of information integration and transmission during early skill learning.

**Significance Statement:** Early learning of moderately difficult skill sequences develops over periods of rest interspersed with practice (micro-offline gains). We demonstrate here a link between alpha and low-beta oscillatory activity in a cingulate-hippocampo-striato network hubs and rest intervals of early learning.

## Introduction

Learning new skills like playing musical instruments or sports is common during performance of activities of daily living (Davare et al., 2007; Friedman and Korman, 2016; Newell et al., 2009; Perez et al., 2008; Song, 2009). Initial practice of a new sequential skill of moderate difficulty results in marked skill improvements that develop predominantly over periods of rest interspersed with practice (micro-offline gains) (Bonstrup et al., 2019; Dayan and Cohen, 2011; Jacobacci et al., 2020; Robertson, 2019). Micro-offline gains in the context of early learning are associated with brain activity and connectivity changes that include wakeful neural replay (Buch et al., 2021) in cortico-subcortical networks engaging the primary motor (Mary et al., 2016; Sugata et al., 2020) and premotor cortices (Wu et al., 2014), the limbic system and hippocampus (Jacobacci et al., 2020), and the basal ganglia (Albouy et al., 2013b; Cataldi et al., 2022).

Network hubs optimize communication and information transfer (Bassett and Bullmore, 2009; Buckner et al., 2009; Hwang et al., 2013; Oldham and Fornito, 2019) supporting different memory-related processes (Bassett and Bullmore, 2009; Hwang et al., 2013; Oldham and Fornito, 2019), attention (Betti et al., 2018), error detection (Buckner et al., 2009; Ide and Li, 2011), and motor planning (Yeom et al., 2020). The connectivity pattern of network hubs informs on the neural dynamics supporting task performance (Crossley et al., 2013; Fair et al., 2009; Yeom et al., 2020). Different brain oscillatory rhythms (Buzsaki and Draguhn, 2004) expressed within these hubs are thought to relay information in the context of different activities: alpha (8–13 Hz, low sensorimotor rhythm linked to motor action) (Spitzer et al., 2014; Sugata et al., 2014), low-beta (13-16Hz, high sensorimotor rhythm also linked to motor action) (Pineda, 2005; Zapala et al., 2015), high-beta (16-30Hz, linked to attentional states) (Susam et al., 2022; Voloh and Womelsdorf, 2018), low-gamma (30-50 Hz, involved in motor processing) (Fernandez-Ruiz et al., 2023) and high-gamma (>60 Hz, linked to memory formation) (Haque et al., 2020).

Magnetoencephalography (MEG) is an ideal technique for investigating brain oscillatory network hubs linked to motor skill learning due to its high temporal resolution and ability to capture neural activity with millisecond-level precision (Baillet, 2017). Skill learning induces fast interactions between brain regions over very short time scales requiring high temporal resolution methods to characterize the neural dynamics (Dayan and Cohen, 2011). Importantly, MEG is sensitive to oscillatory brain activity in both cortical (Baillet, 2017) and subcortical regions including the hippocampus, amygdala, putamen, thalamus (Lopez-Madrona et al., 2022; Pizzo et al., 2019) and striatum (Alberto et al., 2021; Sepe-Forrest et al., 2021)

Here, we used MEG to characterize oscillatory brain activity in principal network hubs contributing to micro-offline gains during rest intervals of early skill learning (Bonstrup et al., 2020; Brooks et al., 2024; Johnson et al., 2023). We identified two distinct groups of network hubs which correlated with micro-offline gains: 1) a group distributed between bilateral anterior cingulate (ACC) and caudate regions characterized by strong alpha band (8-13Hz) interactions; and 2) another group located in bilateral caudate regions and the right putamen characterized by low-beta band (13-16Hz) interactions. Both hub groups showed strong connectivity with the hippocampus, parahippocampal cortex, and lingual and fusiform gyri.

## Materials and methods

### Participants

Thirty-three naïve right-handed healthy volunteers (26.6 ± 0.87 years; 16 female) participated in the study (Bonstrup et al., 2019). Two participants were excluded from the analysis due to technical problems which resulted in incomplete MEG data acquisition. All participants with a normal neurological examination provided written informed consent. The Combined Neuroscience Institutional Review Board of the National Institute of Health (NIH) approved the study protocol. The study sample size was determined via an *a priori* power analysis based on previous skill learning effects observed for the same task (Bonstrup et al., 2019). Active musicians were excluded from the study (Ruiz et al., 2009a; Ruiz et al., 2009b). All data were de-identified and permanently unlinked from all personally identifiable information (PII) before the analysis.

### Task

Participants were asked to perform a 5-item numerical sequential motor task where they repetitively typed the sequence displayed on a projector screen (4-1-3-2-4) as fast and as accurately as possible with their non-dominant, left hand on a response pad (Cedrus LS-LINE, Cedrus Corp). Keypress 1 to 4 corresponded to the little, ring, middle, and index fingers, respectively. Participants performed the task over 36 practice trials lasting 10 sec each (**Fig. 1**). Rest periods of 10 sec were interspersed between practice trials. During practice periods, participants were asked to focus their eyes on the five-item sequence displayed in front of them. During rest periods, participants were instructed to fixate their gaze on a string of five “X” symbols displayed on the monitor. Visual stimuli and keypress response measurements were controlled by a custom script running in E-Prime 2 (Psychology Software Tools, Inc.).

**Fig. 1.**
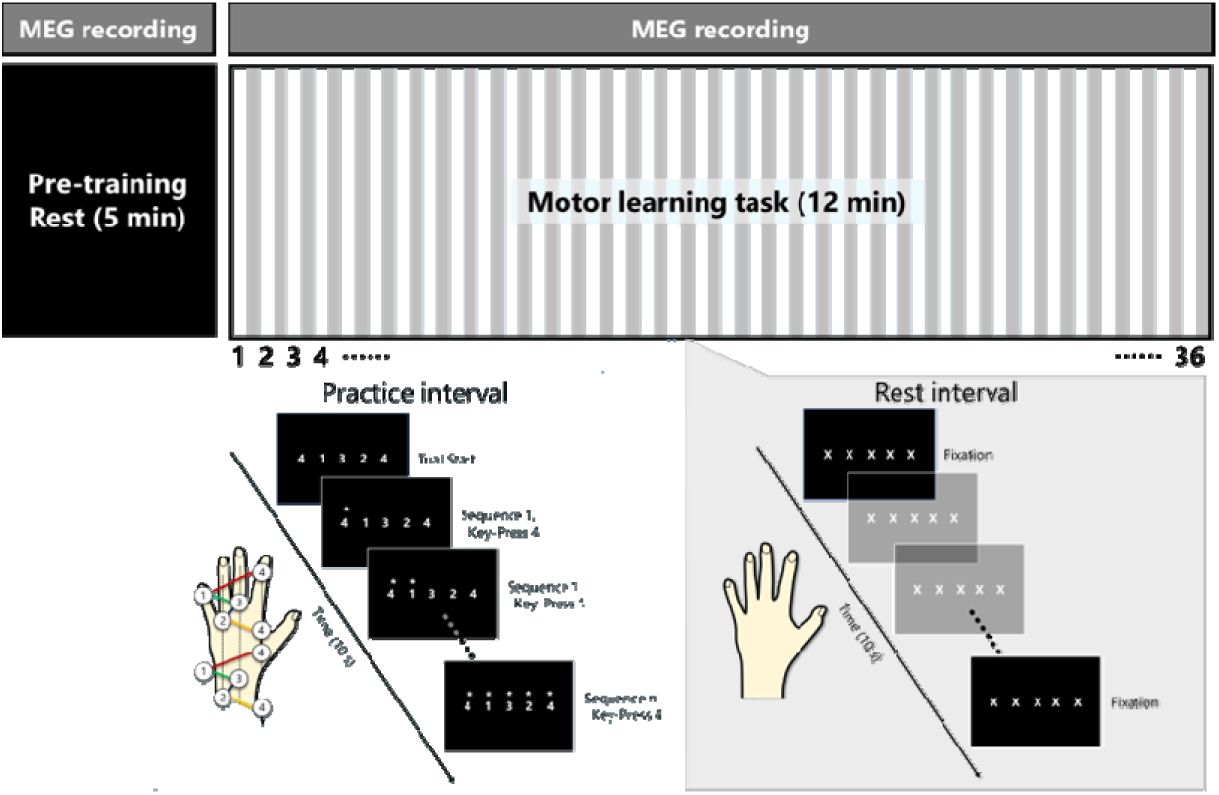
Keypress sequence motor learning task. Subjects were asked to repeatedly type the keypress sequence 41324 with their left non-dominant hand as fast and accurately as possible. After a 5-minute resting MEG data recording (pre-training rest), subjects performed the task over 36 individual trials lasting 10s each (practice intervals). Practice periods were interleaved with 10s waking rest intervals.

### Skill measures

Skill learning behavioral results were reported in Bönstrup et al. (2019). The correct sequence speed was determined by calculating the inverse of the time (in seconds) required to complete a single full iteration of a correct 5-item sequence. Individual keypress responses were considered correct if they matched any possible circular shifts of the 5-item sequence displayed on the monitor (e.g., 41324). Thus, the skill measure was assessed within each practice trial at the resolution of individual keypresses, with skill performance summarized as the mean correct sequence speed across the entire trial. Erroneous keypress actions (< 3%) were excluded from the analysis.

#### Early learning

Early learning was the window of practice trials required to reach 95% of the maximal within-session performance at the group level (*T*). It was calculated by fitting the trial-by-trial group average correct sequence speed data with an exponential model:

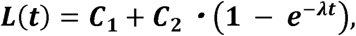

where *L(t)* represented the group average correct sequence speed on practice trial, *t*, *C_1_* and *C_2_* controlled the initial performance and asymptote (i.e., – extent of learning), and λ controlled the learning rate. Parameters C_1_ (boundary constraints = [0,5]), C_2_ ([0,15]) and λ ([0,2]) were previously estimated using a constrained nonlinear least-squares method (MATLAB’s lsqcurvefit, trust-region-reflective algorithm). Micro-offline gains were calculated as the difference in correct sequence speed between the end of one practice period and the beginning of the next, while micro-online gains were the instantaneous difference in correct sequence speed between the beginning and end of a practice trial. We then calculated cumulative micro-offline and micro-online gains over the period of early learning (Bonstrup et al., 2019).

### MRI

Structural MRI scanning was acquired using a 3T MRI scanner (GE Excite HDxt and Siemens Skyra) with a standard 32-channel head coil for each participant to allow volume-based current source estimation from MEG signals.

### MEG

#### Data acquisition

Continuous MEG was recorded in a magnetically shielded room using a CTF 275 MEG system (CTF Systems, Inc., Canada) at a sampling frequency of 600 Hz. During MEG recording, the participants were seated in a comfortable chair. The MEG system was equipped with a head array of 275 radial 1st-order gradiometer / SQUID channels (Vacuumschmelze, Germany). Two of the gradiometers were eliminated from recording due to malfunctioning. One channel was removed after visual inspection of the data indicated the presence of high noise levels for all recordings. Thus, 272 total channels of MEG data were used in subsequent analysis. Synthetic 3rd order gradient balancing was used to remove background noise. A TTL trigger was used to temporally align the behavioral and MEG data. Head position was monitored using three head localization coils attached to the participant’s nasion, left and right pre-auricular locations. These coil positions were also recorded during the session using a stereotactic neuro navigation system (BrainSight, Rogue Research Inc.) to align the MEG data with the individual’s structural MRI.

#### MEG Preprocessing

MEG data were analyzed using Brainstorm software (Tadel et al., 2011). A notch filter eliminated the power line noise (60 Hz). Eye blinking during the experiment was monitored by an eye tracker (EYELINK 1000 PLUS, SR Research). Infomax Independent Component Analysis (ICA) was fit to retain 272 independent components. Spatial topographies and time series of all components were visually inspected and those determined to be strongly associated with known electrocardiogram, eye movements and blinks were removed. Preprocessed MEG data during early learning were visually inspected and we removed rest and practice 10 sec time intervals strongly contaminated by electromyogram and body movements. This procedure removed 20.97 ± 18.7 sec (mean±std.) of MEG data.

#### Source estimation

Each participant’s T1-weighted brain MRI was used to generate an individual head model (volume-level boundary element method, BEM), including three layers (scalp, skull, and brain). Noise covariance was used to estimate source space current distributions during the 10 sec rest intervals, and during practice periods using the minimum norm estimate (MNE) calculated using the Brainstorm’s procedure (Tadel et al., 2019). Noise covariances were calculated from pre-training resting MEG data. The 5-min recordings were divided into 30, 10-s long epochs. All epochs were baseline-corrected by subtracting the mean amplitude from individual channels. Covariance matrices were calculated for individual epochs and then averaged to compute the final noise estimate. Next, whole-brain MNE source estimates were calculated using a 5x 5x 5 mm voxel size. Source data were parcellated into eighty-six brain regions based on the AAL3 atlas (Rolls et al., 2020) and averaged within each brain region (**Fig. 2**, upper left). Next, source MEG data were concatenated separately over rest intervals and practice of early learning. Power spectrum density (PSD) of MEG source data (Welch’s method, (Welch, 1967)) revealed comparable event-related desynchronizations (ERDs, **Fig. S1A**) and synchronizations (ERSs, **Fig. S1B**) in superficial and deep brain regions, consistent with previous reports (Pu et al., 2018; Sepe-Forrest et al., 2021).

**Fig. 2.**
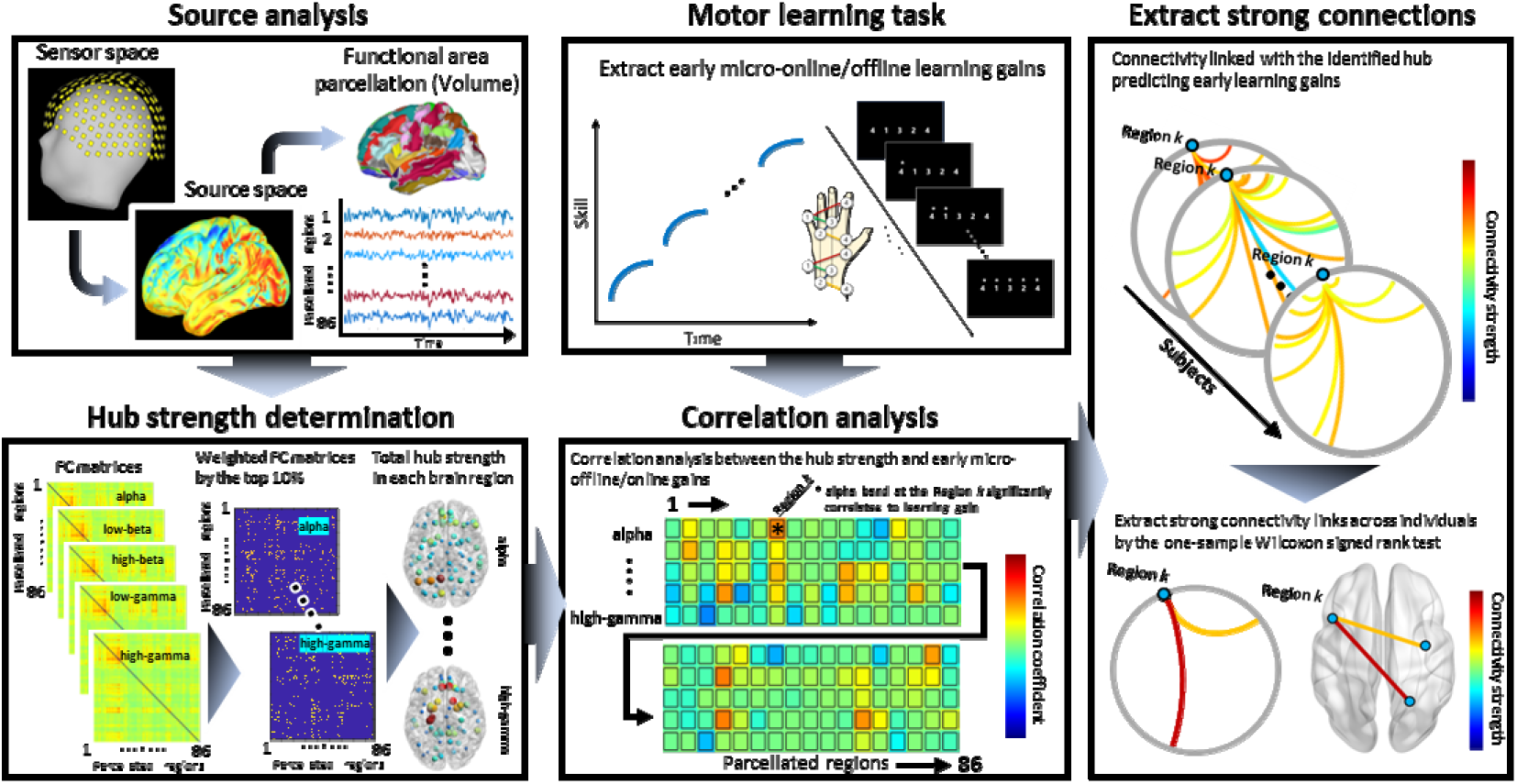
Analysis pipeline. Raw MEG data was pre-processed, source-localized, and parcellated into 86 brain regions using the AAL3 atlas (Rolls et al., 2020). Functional connectivity (FC) analyses were performed using lagged coherence (Pascual-marqui, 2007) in five frequency bands (alpha, low-beta, high-beta, low-gamma, and high-gamma). Hub strengths were determined by the sum of the top 10^th^ reliable connections for each parcellated brain regions and frequency band (Asadi et al., 2021; Liu et al., 2014; Sporns et al., 2021). Next, we measured the correlation coefficient between the hub strength and behavior. Finally, for these ROIs and frequencies associated with learning gains, Bonferroni-corrected one-sample Wilcoxon signed rank tests identified the most relevant connections related to learning.

#### Hub strength determination

In the present study, we calculated the network hub strength for each parcellated brain region to extract crucial network hubs related to skill learning. Hub strength during rest intervals was assessed with micro-offline gains and during practice intervals with micro-online gains.

In concrete, we first calculated functional connectivity between all pairs of parcellated brain regions (86×86) over rest and practice intervals using lagged coherence (1s sliding window length with 80% overlap), an approach that minimize distortions in volume conduction (Pascual-marqui, 2007) (**Fig. 2**, lower left). Next, we identified the top 10% functional connectivity values between these pairs (Asadi et al., 2021; Liu et al., 2014; Sporns et al., 2021). We then calculated the sum of the top 10% functional connectivity values for each of the 86 parcellated brain regions (hub strength) (Hallquist and Hillary, 2019). This procedure was implemented for each of the following frequency bands, chosen to inform on different functional processes and neural computations contributing to brain dynamics (Buzsaki and Draguhn, 2004): alpha (8–13 Hz) (Spitzer et al., 2014; Sugata et al., 2014), low-beta (13-16Hz) (Pineda, 2005; Zapala et al., 2015), high-beta (16-30Hz) (Susam et al., 2022; Voloh and Womelsdorf, 2018), low-gamma (30-50 Hz) (Fernandez-Ruiz et al., 2023)(Fernandez-Ruiz et al., 2023) and high-gamma (70-90 Hz) (Axmacher et al., 2010).

#### Network hubs operating during early skill learning

Next, we identified the hub regions best predictive of skill gains taking place during rest intervals (micro-offline) and practice (micro-online) across all parcellated brain regions and frequency bands (FDR-corrected Pearson’s correlation coefficient (Benjamini and Hochberg, 1995); **Fig. 2**, lower middle). Finally, we identified the connectivity links from each of these hubs stronger than zero-median distribution (Bonferroni-corrected Wilcoxon signed rank tests; **Fig. 2**, right (Aznarez-Sanado et al., 2021; Pahapill et al., 2020)).

## Results

### Behavioral results

Modeling analysis showed that the group average 95% skill was achieved at trial 11 (**Fig. 3A**). Speed of correct sequences during this early learning period increased rapidly by 0.52 ± 0.21 sequence/s (*t* = 13.74, Cohen’s *d* = 2.468, *p* < 0.001; **Fig. 3B, C**). Most skilled gains developed during rest intervals (micro-offline gains: 0.65 ± 0.79 [mean ± SD] sequence/s; two-tailed one-sample t-test: *t* = 4.52, Cohen’s *d* = 0.81, *p* < 0.001) rather than during practice (micro-online gains: −0.12 ± 0.83 sequence/s; *t* = −0.77, Cohen’s *d* = −0.14, *p* = 0.45) periods.

**Fig. 3.**
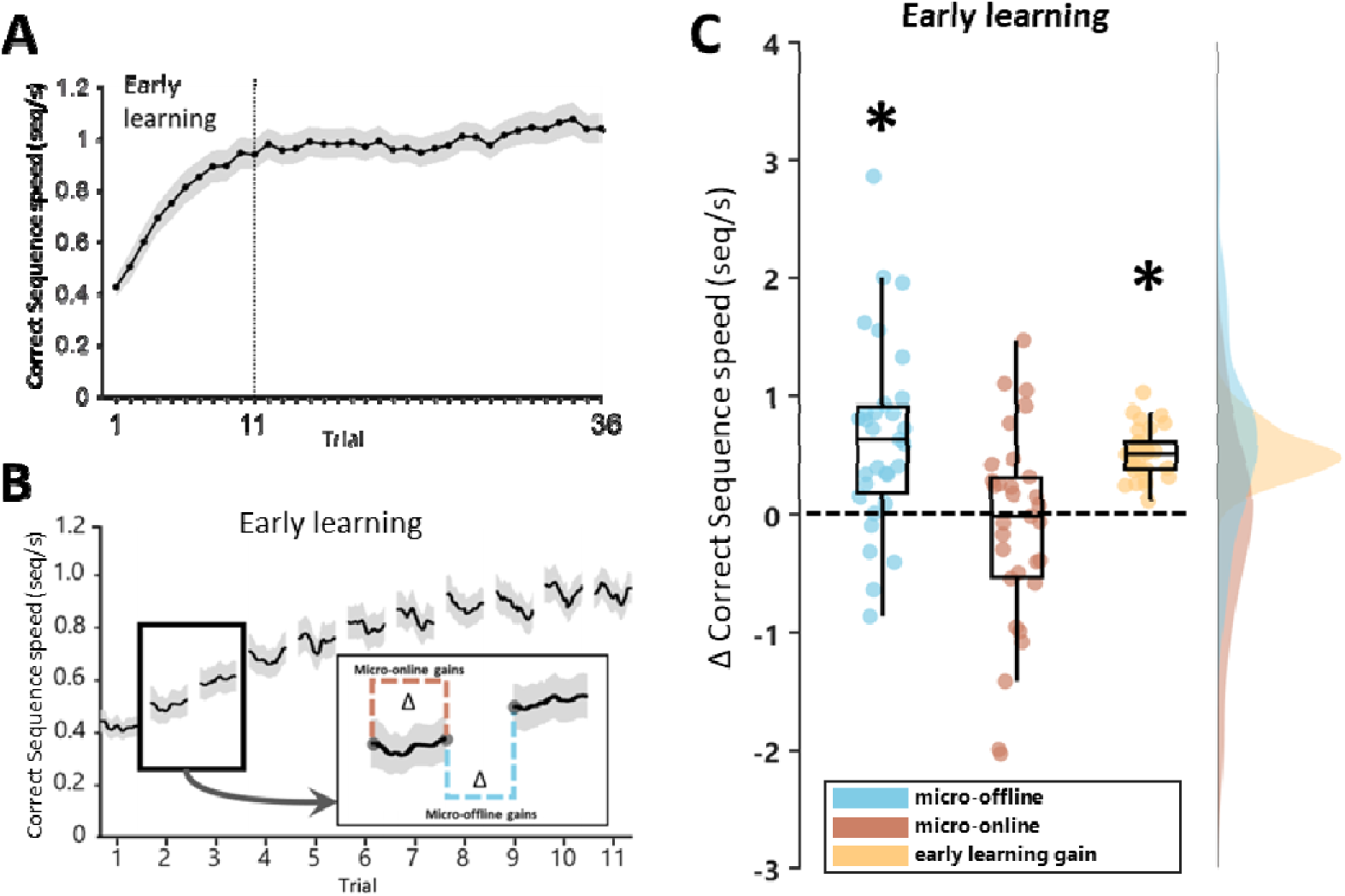
Microscale learning. **(A)** Skill was measured as the correct sequence typing speed (sequence [seq]/s). Mean average performance (mean ± SEM) increased rapidly during early learning (the set of trials required to reach 95% of total learning, in this study trials 1-11, vertical dotted line). **(B)** We then measured performance improvements during (micro-online gains) and in between (micro-offline gains) practice periods of early learning. Micro-online changes were calculated as the difference in tapping speed between the first and the last sequence within each practice period (brown). Micro-offline changes were calculated as the difference between the last sequence within a practice period and the first sequence in the following practice period (blue). **(C)** Significant improvements in skill were observed in micro-offline gains (blue) and early total gain (brown, * p < 0.001, two-tailed one-sample t-test for each training partition, corrected by Bonferroni). The thick horizontal line, the box, and the whiskers represent the median, interquartile range, and maximum and minimum scores, respectively. The curves on the right side depict the data distribution.

*Micro-offline gains correlate with hub strength during rest periods of early learning*. During rest intervals, micro-offline gains correlated with hub strength in the alpha band in the right subgenual ACC (ACC sub) (*r* = 0.568, *p* = 0.00087), pregenual ACC (ACC pre) (*r* = 0.601, *p* = 0.00035), and bilateral supragenual ACC (ACC sup) (left: *r* = 0.652, *p* = 0.000072, right: *r* = 0.570, *p* = 0.00080) and caudate nuclei (left; *r* = 0.588, *p* = 0.00061, right; *r* = 0.563, *p* = 0.00097, **Fig. 4 and Fig. S2**) and in the low-beta band, in the right putamen (*r* = 0.582, *p* = 0.00059) and bilateral caudate nuclei (left; *r* = 0.565, *p* = 0.00093, right; *r* = 0.596, *p* = 0.00040, **Fig. 4** and **Fig. S2**). Comparable results were obtained with a correlation analysis robust against outliers (Spearman’s correlation, **Fig. S2**). During practice intervals, micro-online skill gains did not correlate with any regional or oscillatory hub strength.

**Fig. 4.**
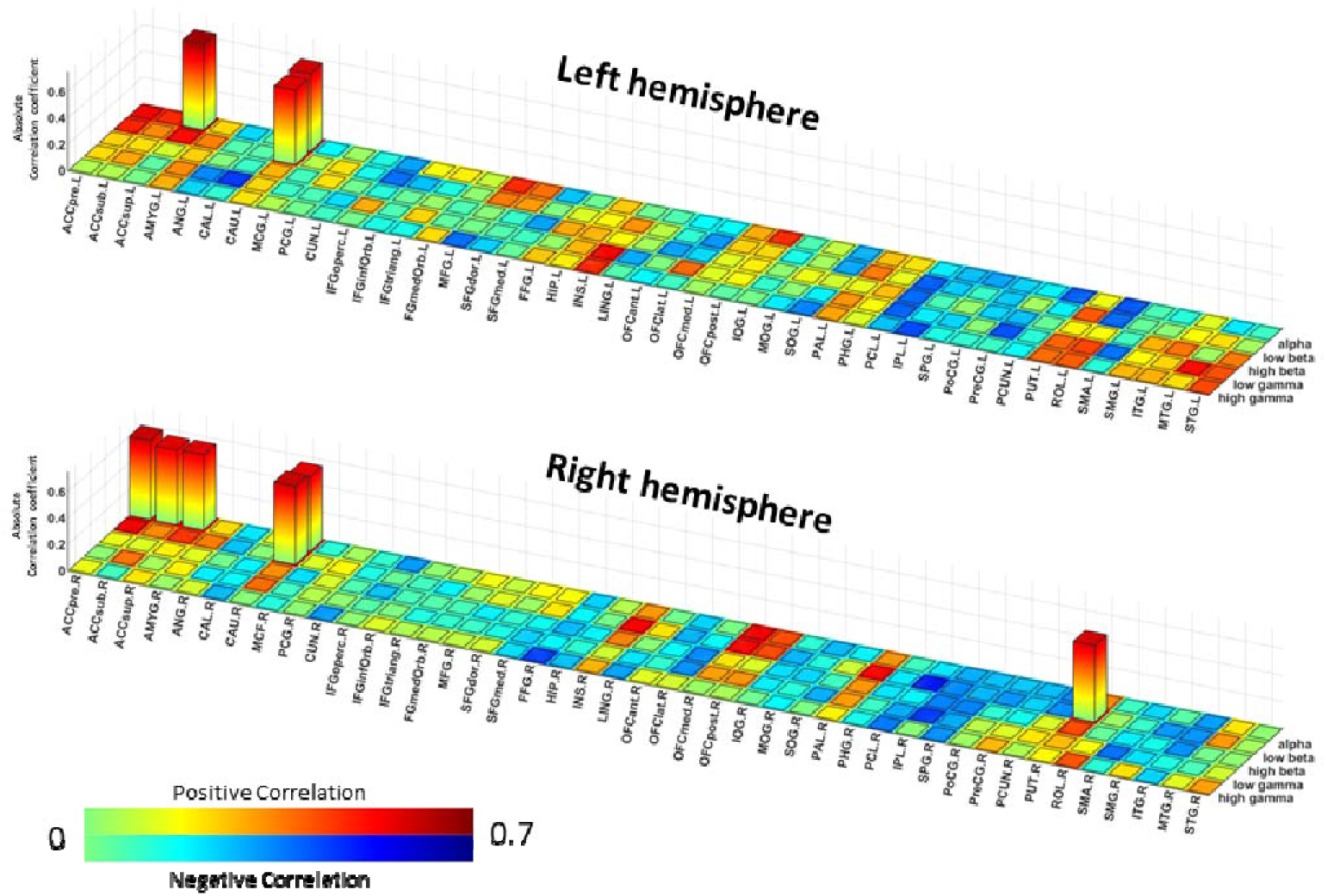
Hub strength during early learning correlated with micro-offline gains. Hub strength in bilateral ACCs and bilateral caudate regions in the alpha band and two bilateral caudate and right putamen regions in the low-beta band significantly correlated with micro-offline gains. Heat maps depict correlation values. Only significant parcellated brain regions are shown by the bar plots. We did not find significant negative correlations. All p-values were corrected by False Discovery Rate (FDR) with a q-value of 0.05.

Next, we identified the strongest connections linked with micro-offline network hubs identified in the previous paragraph (ROI-frequency band combinations, **Fig. 4**). In the alpha band, three of the six ROIs depicted in **Fig. 4** showed significantly strong connectivity links (the right ACC sub and bilateral caudate nuclei, **Fig. 5A**): the right ACC sub with the left hippocampus (0.204 ± 0.045, *p* = 0.042), parahippocampal cortex (0.215 ± 0.045, *p* = 0.041), and amygdala (0.2114 ± 0.039, *p* = 0.042; **Fig. 5A,B** left); the left caudate nucleus with the left inferior temporal gyrus (ITG) (0.177 ± 0.075, *p* = 0.041), fusiform gyrus (0.122 ± 0.095, *p* = 0.021), posterior cingulate cortex (PCC) (0.114 ± 0.097, p = 0.041), and the right lingual gyrus (0.111 ± 0.094, *p* = 0.042); the right caudate nucleus with the left hippocampus (0.124 ± 0.106, *p* = 0.042) (**Fig. 5A,B** right).

**Fig. 5.**
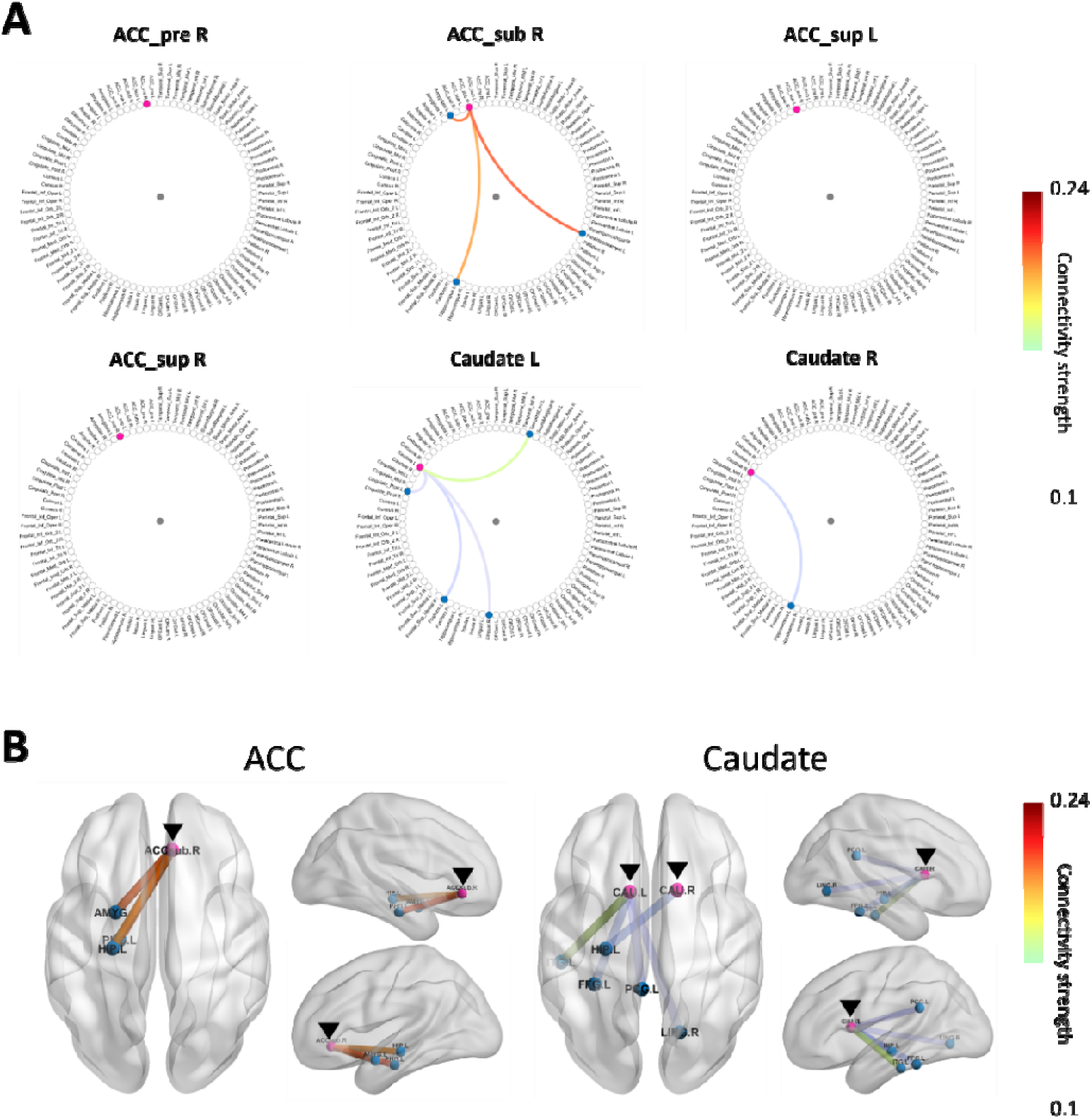
Connectivity links in the alpha band. **A**. Circular plots depict the alpha band connectivity links from each of the six regions identified in Figure 4. Significant (Bonferroni-corrected) connectivity links emerged from three of the six regions (right subgenual ACC or ACC sub and bilateral caudate) communicating the right ACC with the left hippocampus, parahippocampal cortex, and amygdala; the left caudate nucleus with the left inferior temporal gyrus, fusiform gyrus, posterior cingulate cortex (PCC), and right lingual cortex; and the right caudate nucleus with the left hippocampus. **B**. Cartoons depict these significant connectivity links. Pink circles with black arrowheads indicate ROIs significantly correlated with early micro-offline gains. Colored bold lines between pink and blue circles illustrate connectivity strength.

In the low-beta band, all three ROIs identified in **Fig. 4** showed significantly strong connectivity links (right putamen and bilateral caudate nuclei, **Fig. 6A**) the right putamen with the left hippocampus (0.223 ± 0.064, *p* = 0.010), parahippocampal cortex (0.234 ± 0.069, *p* = 0.0052), amygdala (0.214 ± 0067, *p* = 0.039), PCC (0.203 ± 0.049, *p* = 0.020), fusiform gyrus (0.228 ± 0.055, *p* = 0.020), and ITG (0.233 ± 0.065, *p* = 0.020), the right PCC (0.215 ± 0.052, *p* = 0.039), fusiform gyrus (0.195 ± 0.038, *p* = 0.039), and lingual gyrus (0.202 ± 0.061, *p* = 0.020; **Fig. 6A,B** left); the left caudate nucleus with the left hippocampus (0.170 ± 0.107, *p* = 0.010), parahippocampal cortex (0.217 ± 0.071, *p* = 0.0026), PCC (0.140 ± 0.106, *p* = 0.010), fusiform gyrus (0.139 ± 0.113, *p* = 0.020), medial orbitofrontal cortex (OFC med) (0.201 ± 0.107, *p* = 0.020), the right fusiform gyrus (0.150 ± 0.107, *p* = 0.02), and the lingual gyrus (0.138 ± 0.108, *p* = 0.020); the right caudate nucleus with the left hippocampus (0.172 ± 0.105, *p* = 0.0052), parahippocampal cortex (0.219 ± 0.090, *p* = 0.0026), amygdala (0.212 ± 0.080, *p* = 0.0051), fusiform gyrus (0.181 ± 0.107, *p* = 0.020), right fusiform gyrus (0.141 ± 0.091, *p* = 0.020) and lingual gyrus (0.168 ± 0.089, *p* = 0.0051, **Fig. 6A, B** right).

**Fig. 6.**
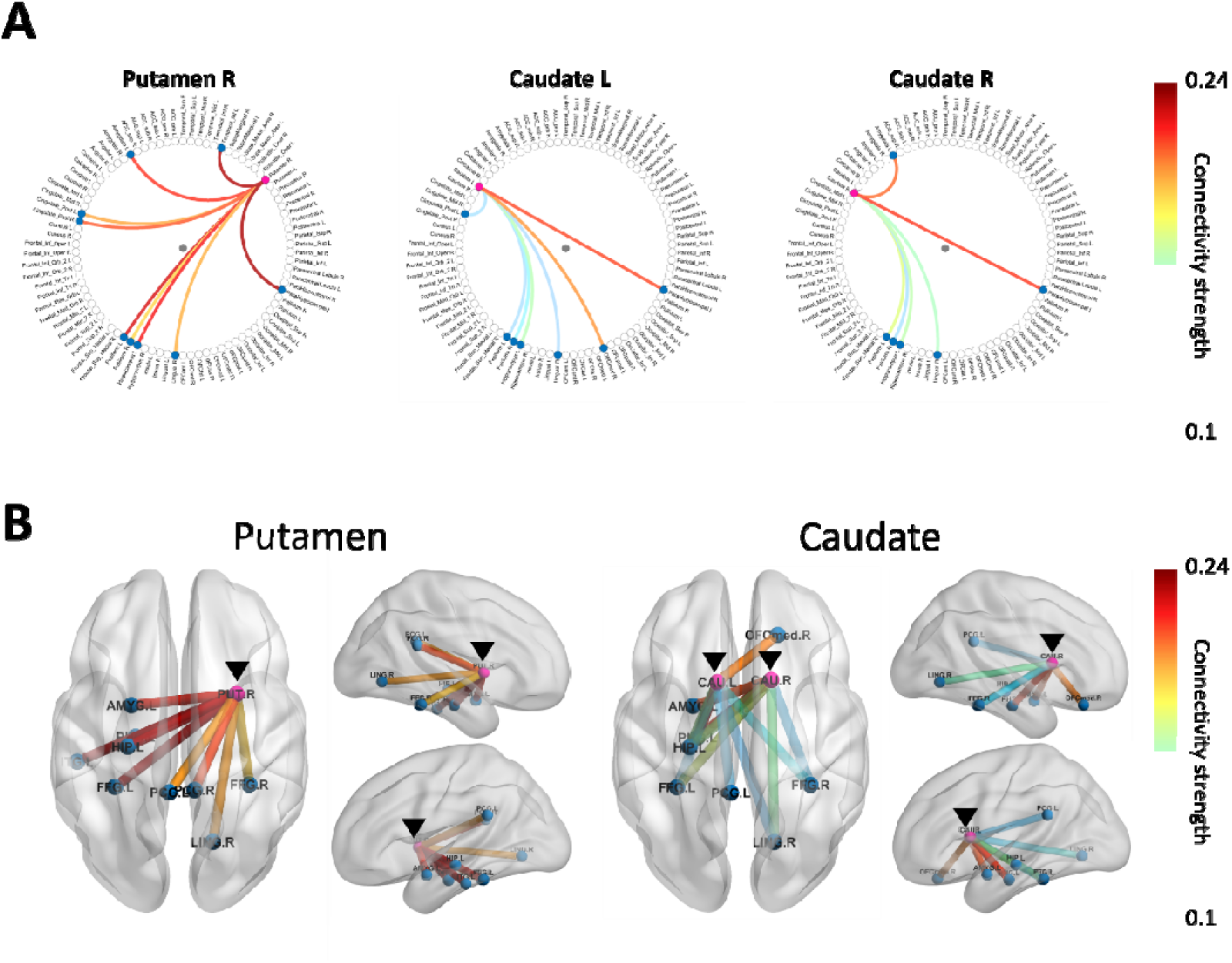
Connectivity links in the low-beta band. **A.** Circular plots depict low-beta band connectivity links for each of the three regions identified in Figure 4. Significant (Bonferroni-corrected) connectivity links emerged from all three regions (bilateral caudate and right putamen) communicating the right putamen with the left hippocampus, parahippocampal cortex, amygdala, inferior temporal gyrus, right lingual gyrus, bilateral PCC, and fusiform gyri; the left caudate nucleus with the left hippocampus, parahippocampal cortex, and PCC, the right medial orbitofrontal cortex (OFCmed), lingual gyrus, and the bilateral fusiform gyri; and the right caudate nucleus with the left hippocampus, hippocampal cortex, amygdala, the right lingual gyrus, and the bilateral fusiform gyri. **B**. Cartoons depict topographically the significant connectivity links. Pink circles with black arrowheads indicate ROIs significantly correlated with early micro-offline gains. Colored bold lines between pink and blue circles illustrate connectivity strength.

## Discussion

Performance improvements during skill learning develop over periods of practice and rest (Walker et al., 2003). Recent work showed that early learning of moderately complex sequential skills develops predominantly during rest rather than during practice intervals (i.e. – expressed as micro-offline gains) (Bonstrup et al., 2019). Two non-mutually exclusive mechanisms have been posited to contribute to micro-offline gains: (1) rapid consolidation of skill (Bonstrup et al., 2019) or (2) recovery from fatigue (Gupta and Rickard, 2022; Hull, 1943). Experimentally shortening practice duration—thus reducing fatigue—does not modify micro-offline gains, which is inconsistent with latent motor slowing resulting from within-practice performance decrements (Bonstrup et al., 2020). On the other hand, the motor memory stabilizes during rest intervals of early learning (Bonstrup et al., 2020) (Bonstrup et al., 2020) supporting the view that micro-offline gains in this context are expressions of a form of rapid consolidation(Brooks et al., 2024; Jacobacci et al., 2020; Robertson, 2019). One possible mechanism supporting micro-offline gains is hippocampo-sensorimotor neural replay (Buch et al., 2021; Jacobacci et al., 2020). The hippocampus and the sensorimotor cortex are active contributors to the network involved in motor learning, which also includes the basal ganglia (Albouy et al., 2013a; Albouy et al., 2013c) and anterior cingulate (Cataldi et al., 2022). We now identify the topographical hubs within this network and their prominent oscillatory frequencies that operate during rest intervals in which micro-offline gains of early learning occur.

We found that the strongest network hubs during rest intervals of early learning were the anterior cingulate, the caudate and the putamen. The anterior cingulate (ACC) receives projections from the hippocampus and amygdala (Stevens et al., 2011) and has strong connections with the sensorimotor divisions of the basal ganglia and the thalamus (Doyon et al., 2009; Miyachi et al., 1997). ACC activity is modulated by early motor learning (Berlot et al., 2020; Doyon and Benali, 2005; Gonzalez and Burke, 2018; Iwane et al., 2023) and encoding of the serial order of sequence components (Procyk et al., 2000) (Procyk et al., 2000). It is also thought to integrate motor planning with cognitive and emotional feedback, particularly during rest (Gusnard et al., 2001).

The identification of striatal hubs is in line with previous reports linking dorsal striatal function with learning (Cataldi et al., 2022; Fitzroy et al., 2021; Gusnard et al., 2001; Miyachi et al., 1997; Wessel et al., 2023; Yin et al., 2009), particularly for motor sequences (Andersen et al., 2020; Choi et al., 2020; Doyon and Benali, 2005; Hikosaka et al., 1999; Janacsek et al., 2020). Dorsal striatal activity also contributes to memory consolidation (Albouy et al., 2008; Debas et al., 2010; Peigneux et al., 2003). Specifically, the caudate nucleus receives afferents from the cerebral cortex, midbrain and thalamus, is active during sequence learning (Maquet et al., 2000) in association with performance of reproducible (less variable) motor behavior (Albouy et al., 2012) and is engaged in action selection, reinforcement, and updating of motor planning (Hikosaka et al., 2002; Miyachi et al., 2002). The putamen is an input nucleus to the basal ganglia, has an anterior associative subregion active during early skill learning as well as a posteroventral sensorimotor region more involved in later learning stages (Lehericy et al., 2005).

The main connectivity links from the ACC, caudate and putamen hubs observed in the present study are with the hippocampus, parahippocampal cortex, and lingual and fusiform gyri. The hippocampus is engaged in memory processing (Squire, 1982, 1986) and rapid wakeful consolidation in the context of early skill learning (Buch et al., 2021; Gann et al., 2021; Jacobacci et al., 2020; Mylonas et al., 2024) as well as sleep-dependent consolidation occurring over several hours (Albouy et al., 2013b; Lewis et al., 2018; Liu et al., 2019).

Hubs identified during rest intervals of early learning operated within alpha and low beta brain oscillatory frequencies previously linked to sensorimotor processes (Pineda, 2005; Zapala et al., 2015). Modulation of alpha activity is associated with attention (Basar et al., 2001; Palva and Palva, 2007), motor execution (Leocani et al., 2001; Manuel et al., 2018; Salmelin and Hari, 1994; Wang et al., 2022) and hippocampo-striatal binding contributing to memory processing (Herweg et al., 2016). Modulation of beta activity is observed during motor action (Barone and Rossiter, 2021; Schmidt et al., 2019), rapid consolidation of skill (Bonstrup et al., 2019; Pollok et al., 2014), and NMDA-dependent synaptic plasticity (Wischnewski et al., 2019) particularly in the striatum, where it is associated with skill acquisition (Koralek et al., 2012). Overall, these findings are consistent with the involvement of sensorimotor rhythms in motor preparation, sensorimotor integration, and attention—all of which are active processes during rest intervals which could support micro-offline gains (Engel and Fries, 2010; Ramos-Murguialday et al., 2013).

In summary, we report that micro-offline gains—which contribute to virtually all total early learning of a moderately difficult skill—is associated with strong cingulate-hippocampo-striato connectivity hubs in the alpha and low-beta oscillatory range. These connectivity hubs may contribute to the binding of individual sequence action components of complex human motor sequences during early learning rest intervals.

## Acknowledgments

We thank Tasneem Malik and Michele Richman for their support. This work utilized the NIMH MEG Core Facility, NIH Functional Magnetic Resonance Imaging Core (FMRIF) Facility, and the computational resources of the NIH HPC Biowulf cluster (http://hpc.nih.gov).

