## Supplemental Figures for "Cingulate and striatal hubs are linked to early skill learning"

**Abbreviated title**: Striatum network for micro-offline learning

Hisato Sugata [1,2,4,5], Fumiaki Iwane [1,4], William Hayward [1], Valentina Azzollini [1], Debadatta Dash [1], Roberto F Salamanca-Giron [1], Marlene Bönstrup [3], Ethan R Buch [1], Leonardo G Cohen [1,5]

**Affiliations**
[1] Human Cortical Physiology and Neurorehabilitation Section, NINDS, NIH, Bethesda, MD, USA
[2] Faculty of Welfare and Health Science, Oita University, Oita, Japan
[3] Department of Neurology, University of Leipzig Medical Center, 04103, Leipzig, Germany
[4] Equal Contribution
[5] Lead Contact

Contents:
Supplementary Figures 2

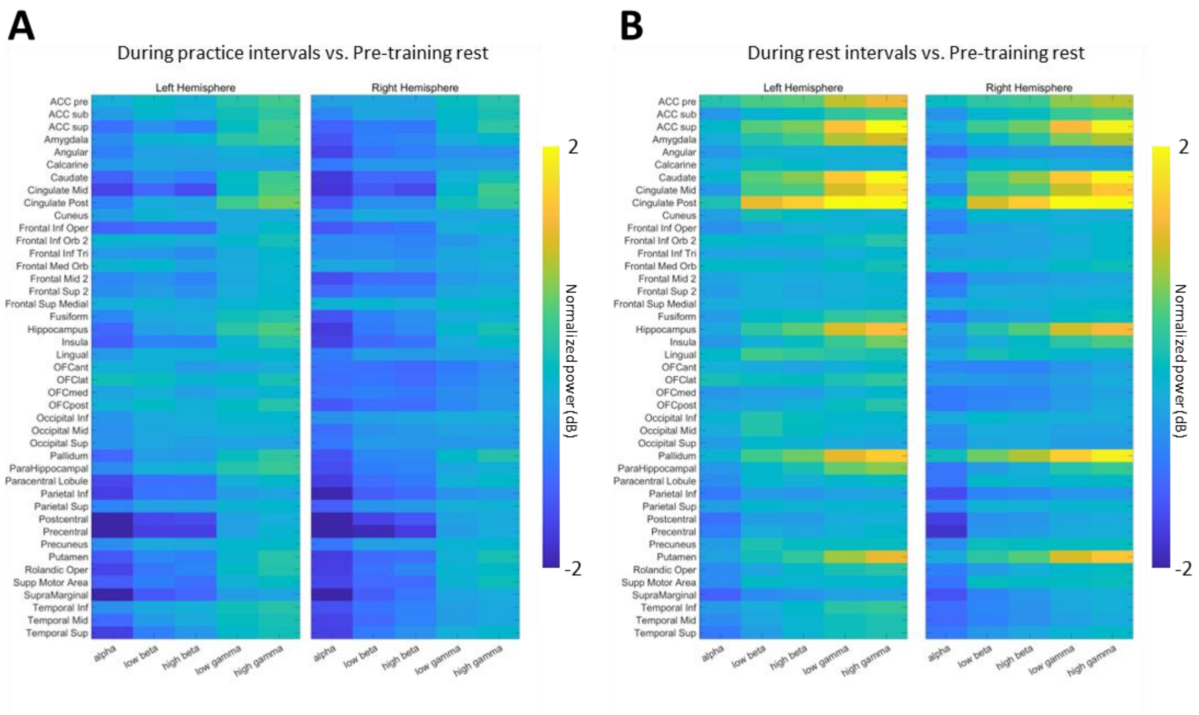


**Fig. S1. Results of power spectrum analysis.**

**A.** In the practice intervals, event-related desynchronizations (ERDs) were induced in deep brain structures, including the caudate nucleus, putamen, and hippocampus, compared to pre-training rest.

**B.** In the rest intervals, event-related synchronizations (ERSs) were observed in similar deep brain regions compared to pre-training rest.

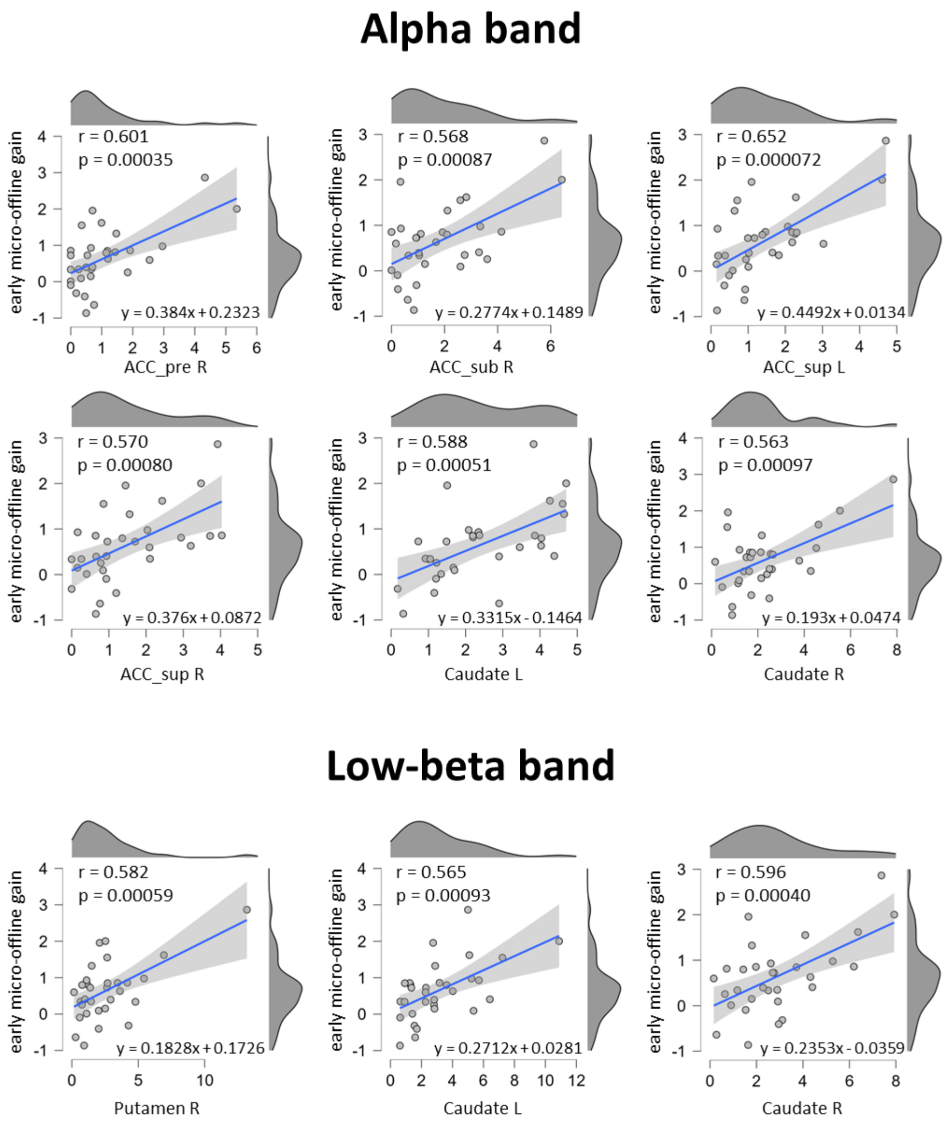


**Fig. S2. Scatter plots depicting the relationship between hub strength and micro-offline gains in parcellated brain regions identified in Figure 4 in alpha and low-beta bands in all subjects.**

Shaded areas show 95% confidence intervals. The x-axis shows hub strength from each significant region to the rest of the brain (calculated by summing up the connectivity values projected from all other parcellated brain regions). The y-axis indicates early micro-offline gains. All FDR-corrected p-values were displayed in the figure. Comparable results were obtained using an approach robust against outliers (Spearman's correlation) (Alpha band: ACC_pre R, ρ = 0.527, p = 0.002, ACC_sub R, ρ = 0.434, p = 0.015, ACC_sup L, ρ = 0.582, p = 0.0007, ACC_sup R, ρ = 0.585, p = 0.0005, Caudate L, ρ = 0.627, p = 0.0002, Caudate R, ρ = 0.532, p = 0.0024, Low-beta band: Putamen R, ρ = 0.367, p = 0.043, Caudate L, ρ = 0.477, p = 0.0076, Caudate R, ρ = 0.457, p = 0.011).
